# Species specificity and intraspecific variation in the chemical profiles of *Heliconius* butterflies across a large geographic range

**DOI:** 10.1101/573469

**Authors:** Kathy Darragh, Gabriela Montejo-Kovacevich, Krzysztof M. Kozak, Colin R. Morrison, Clarisse M. E. Figueiredo, Jonathan S. Ready, Camilo Salazar, Mauricio Linares, Kelsey J. R. P. Byers, Richard M. Merrill, W. Owen McMillan, Stefan Schulz, Chris D. Jiggins

**Author notes:** Corresponding author Kathy Darragh, Department of Zoology, Downing Street, Cambridge, CB23EJ.

## Abstract

In many animals, mate choice is important for the maintenance of reproductive isolation between species. Traits important for mate choice and behavioural isolation are predicted to be under strong stabilising selection within species, however such traits can also exhibit variation at the population level driven by neutral and adaptive evolutionary processes. Here, we describe patterns of divergence among androconial and genital chemical profiles at inter- and intra-specific levels in mimetic *Heliconius* butterflies. Most variation in chemical bouquets was found between species, but there were also quantitative differences at the population level. We found a strong correlation between interspecific chemical and genetic divergence, however, this correlation varied in intraspecific comparisons. We identified ‘indicator’ compounds characteristic of particular species that included known biologically active compounds, suggesting an approach for identification of candidate compounds for future behavioural studies in novel systems. Overall, the strong signal of species identity suggests a role for these compounds in species recognition, but with additional potentially neutral variation at the population level.

## Introduction

Reproductive isolation between lineages is important for the maintenance of species diversity (Coyne & Orr, 2004). In many animals, mate choice provides a strong pre-mating barrier, maintaining reproductive isolation (Friberg et al., 2008; Gray & Cade, 2000; Martin & Mendelson, 2016; Nagel & Schluter, 1998; Ready et al., 2006; Seehausen et al., 2008; Selz, Pierotti, Maan, Schmid, & Seehausen, 2014). Closely related species often differ in traits important for mate choice, with individuals displaying a preference for conspecific phenotypes (Jiggins, Naisbit, Coe, & Mallet, 2001; Mas & Jallon, 2005; M. J. Ryan & Guerra, 2014; Saveer et al., 2014; Yildizhan et al., 2009). These traits are predicted to show strong species-specific differences (Gerhardt, 1982), and typically should be subject to stabilising selection which can act to decrease intraspecific phenotypic variation (Butlin, Hewitt, & Webb, 1985; Pfennig, 1998; Ptacek, 2000). As a consequence, we would expect to find little trait variability, or at least certain features to be invariant, across species geographic ranges (Benedict & Bowie, 2009; Ferreira & Ferguson, 2002; McPeek, Symes, Zong, & McPeek, 2011; Weber, Mitko, Eltz, & Ramírez, 2016). However, these traits can also exhibit variation both within and between populations of the same species, either due to genetic drift and/or varying selective regimes across their ranges (Bolnick & Kirkpatrick, 2012; M. J. Ryan & Guerra, 2014; M. J. Ryan & Rand, 1993; M. J. Ryan, Rand, & Weigt, 1996).

Signals important for behavioural isolation could arise from the divergence of traits used in intraspecific communication between populations (Johansson & Jones, 2007; Mendelson & Shaw, 2012; M. J. Ryan & Rand, 1993; Smadja & Butlin, 2008). Signal divergence can be driven by various factors, both neutral and adaptive, usually involving multiple evolutionary forces (Leonhardt, Rasmussen, & Schmitt, 2013; Sun et al., 2013). A positive correlation between genetic distance and phenotypic variation is consistent with stochastic processes, such as genetic drift, playing a prominent role (Irwin, Thimgan, & Irwin, 2008). In contrast, a lack of correlation between phenotypic and genetic divergence suggests that selection is shaping the phenotypic variation, perhaps driving divergence in different directions in each population (Campbell et al., 2010; Conrad, Paxton, Assum, & Ayasse, 2018; Hankison & Ptacek, 2008; Mullen, Vignieri, Gore, & Hoekstra, 2009).

Chemical compounds, such as sex pheromones, mediate intraspecific communication in many systems (Wyatt, 2003, 2014). The role of chemical signalling in behavioural isolation is also well established, especially among moth species (Löfstedt, 1993; Smadja & Butlin, 2008). Due to the coordination between detection and production, these pheromone blends are traditionally regarded as being under stabilising selection towards a species stereotype (Löfstedt 1993). Even when species-specific characteristics are present, chemical compounds can exhibit intraspecific variation, with both qualitative and quantitative differences found across a species range (Conrad et al., 2018; Gemeno, Lutfallah, & Haynes, 2000; Groot et al., 2009; Huang et al., 1998; McElfresh & Millar, 2001; M. A. Ryan, Moore, & Walter, 1995; Weber et al., 2016).

Studies of *Heliconius* butterflies have contributed to our understanding of adaptation and speciation (Jiggins, 2008, 2017; Merrill et al., 2015). Despite the reliance of this group on visual cues for mating (Bybee et al., 2012; Finkbeiner, Fishman, Osorio, & Briscoe, 2017; Jiggins et al., 2001; Merrill, Chia, & Nadeau, 2014; Sánchez et al., 2015), it has long been suggested that male pheromones also play a role in pre-mating barriers (Jiggins, 2008; Merrill et al., 2015), but so far only a few species have been studied. Behavioural experiments reveal that chemical signalling in *Heliconius erato, H. melpomene* and *H. timareta* is important for female mate choice (Darragh et al., 2017; Mérot, Frérot, Leppik, & Joron, 2015). Electroantennographic studies have shown that *Heliconius cydno* and *H. melpomene* respond to both con- and heterospecific androconial chemical bouquets (Byers et al., 2019), and have identified individual compounds that are biologically active. Furthermore, studies of *H. cydno*, *H. doris*, *H. hecale*, *H. ismenius*, *H. melpomene*, *H. pardalinus*, *H. sara* and *H. timareta* from Panama, Colombia and Ecuador found that major compounds differ between species (Mann et al., 2017; Mérot et al., 2015), suggesting a potential role in reproductive isolation.

The role of chemical signalling is likely to be especially important in co-mimics, where visual signals alone are not sufficient to identify conspecifics (C. Estrada & Jiggins, 2008; Giraldo, Salazar, Jiggins, Bermingham, & Linares, 2008; Mérot et al., 2013; Sánchez et al., 2015). In addition, chemical compounds could be part of a multimodal aposematic warning signal (Rojas et al., 2018; Rothschild, 1961), with some tentative evidence that co-mimics exhibit similar chemical bouquets to aid recognition by predators (Mann et al., 2017).

Here, we describe the chemical profiles of seven species of *Heliconius* from over 250 individuals collected across the Neotropics. We focus on the co-mimetic species *H. melpomene* and *H. erato* that are distributed widely across the Neotropics and analysed both wing androconial and genital compounds of male butterflies. It would be challenging to conduct electrophysiological experiments on large numbers of species and populations, and therefore identifying the biologically active components in pheromone blends across a radiation is beyond the scope of a single study. Instead, we hypothesise that biologically relevant signals for species recognition are likely to be under stronger stabilising selection compared to other compounds, and therefore the consistency of phenotypic variation across the geographic range of a species might provide an initial approach to detect putatively biologically active compounds important for mate choice. We use *H. melpomene* as a test-species due to the availability of behavioural and electrophysiological data to investigate this approach, by evaluating consistency in compound blends across different localities.

The extensive dataset analysed here allows us to test evolutionary hypotheses, as well as identifying interesting candidate compounds for future behavioural studies. Firstly, we investigated intraspecific variation in chemical profiles of *H. melpomene* and *H. erato* and correlated chemical profile data with both geographic and genetic distances between different localities. Furthermore, to test if the chemical compounds are part of the aposematic co-mimicry signal we sampled two different mimicry rings in western Ecuador and Panama.

## Materials and Methods

### Sampling

Between February 2016 and August 2017 wild males of *Heliconius cydno*, *H. elevatus*, *H. eleuchia*, *H. erato*, *H. melpomene*, *H. sapho* and *H. timareta* were collected from twelve localities. Between two and fifteen males were chemically analysed per population (Fig. 1, Table S1). We follow the latest *Heliconius* taxonomy (Lamas & Jiggins, 2017).

**Figure 1.**
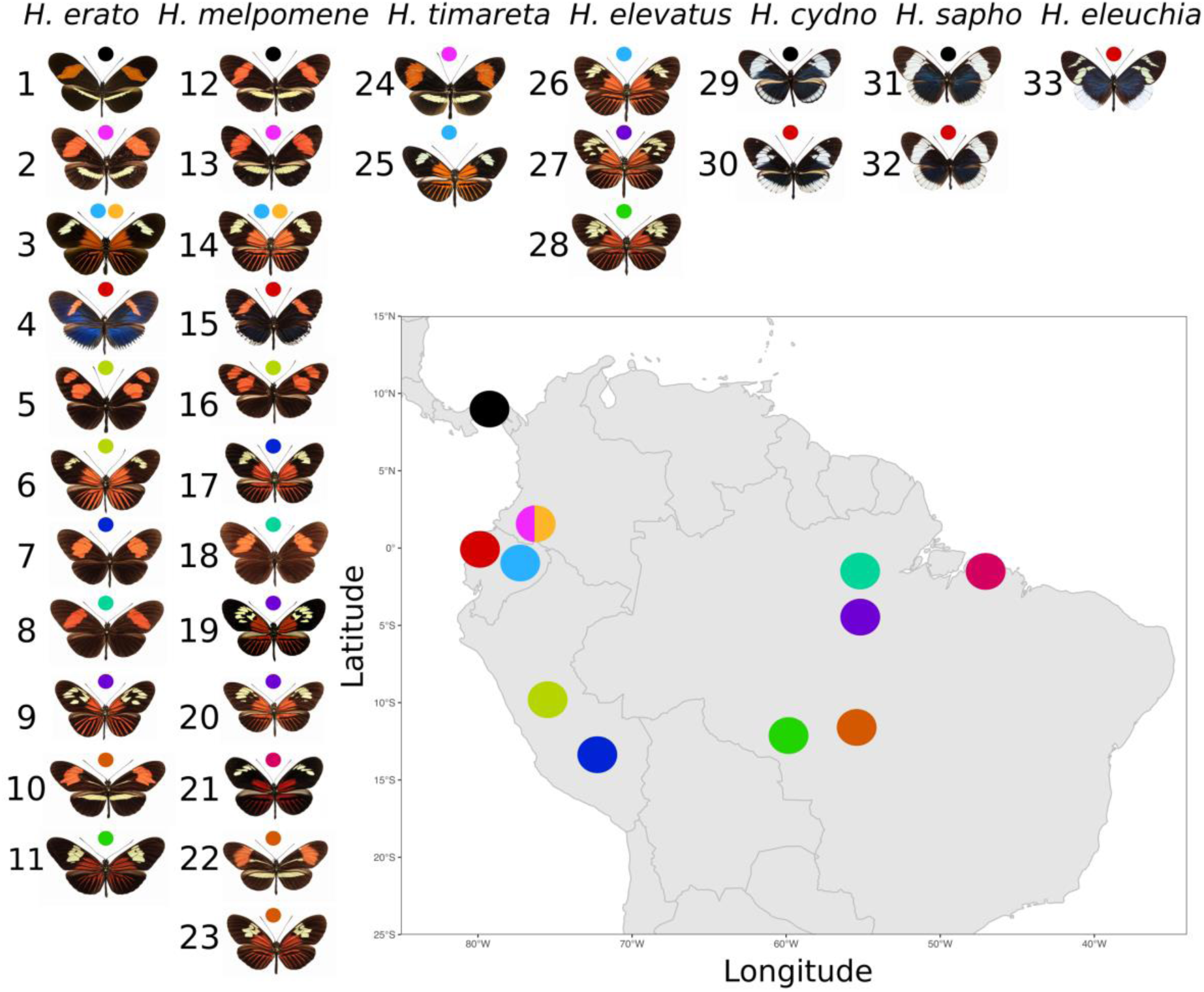
Map indicating species and populations from twelve localities across the Neotropics. H. erato (1-11) 1) *H. e. demophoon* 2) *H. e. dignus* 3*) H. e. lativitta* 4) *H. e. cyrbia* 5) *H. e. microclea* 6) *H. e. emma* 7) *H. e. amphitrite* 8) *H. e. hydara* 9) *H. e. amazona* 10) *H. e. phyllis* 11) *H. e. venustus*. *H. melpomene* (12-23) 12) *H. m. rosina* 13) *H. m. bellula* 14) *H. m malleti* 15) *H. m cythera* 16) *H. m. xenoclea* 17) *H. m. schunkei* 18) *H. m. melpomene* 19) *H. m. madeira* 20) *H. m. thelxiope* 21) *H. m. intersectus* 22) *H. m. penelope* 23) *H. m. burchelli*. *H. timareta* (24-25) 24) *H. t. tristero* 25) *H. t. nov. spp. H. elevatus* (26-28) 26) *H. e. willmotti* 27) *H. e. schmassmanni* 28) *H. e. perchlora*. *H. cydno* (29-30) 29) *H. c. chioneus* 30) *H. c. alithea*. *H. sapho* (31-32) 31) *H. s. sapho* 32) *H. s. candidus*. *H. eleuchia* 33) *H. e. primularis*. See Table S1 for sample numbers.

### Extraction and chemical analysis of tissues

The androconial region of the wing, previously described as the grey-brown overlapping region of the hindwing (Darragh et al., 2017), as well as the genitalia, were dissected for analysis immediately after collection. To extract pheromones, the tissue was soaked in 200μl dichloromethane containing 200ng 2-tetradecyl acetate (internal standard) in 2ml glass vials with PTFE-coated caps (Agilent, Santa Clara, USA) for one hour. The solvent was then transferred to new vials, maintained cool in the field and stored at −20°C upon return. Androconial samples were evaporated to a reduced volume at room temperature prior to analysis. Pheromone extracts were analysed by GC/MS using an Agilent model 5977 mass-selective detector connected to an Agilent GC model 7890B and equipped with an Agilent ALS 7693 autosampler. HP-5MS fused silica capillary columns (Agilent, Santa Clara, USA, 30 m × 0.25 mm, 0.25 µm) were used. Injection was performed in splitless mode (250°C injector temperature) with helium as the carrier gas (constant flow of 1.2 ml/min). The temperature programme started at 50°C, was held for 5 minutes, and then rose at a rate of 5°C/minute to 320°C, before being held at 320°C for 5 minutes. Components were identified by comparison of mass spectra and gas chromatographic retention index with those of authentic reference samples and also by analysis of mass spectra. Components were quantified using 2-tetradecyl acetate as an internal standard. Only compounds eluting earlier than hexacosane were analysed in androconial samples, and those earlier than nonacosane in genital samples (Darragh et al. 2017). We removed compounds that were not found in at least half of all individuals from a given population.

### Calculation of genetic and geographic distance matrices

To explore genetic distance among the studied *H. erato* and *H. melpomene* subspecies, we computed genetic covariance matrices and performed MDS for each species separately. A whole-genome sequence from a representative individual from each subspecies was used (Table S2). Genotypes were inferred from reads mapped to the *H. melpomene* (v2.5) and *H. erato demophoon* genome scaffolds (Challis, Kumar, Dasmahapatra, Jiggins, & Blaxter, 2016; Davey et al., 2017; Heliconius Genome Consortium, 2012; Van Belleghem et al., 2017) with bwa v0.7.15 (Li & Durbin, 2009). We computed a pairwise identical by state (IBS) matrix with a random sampled read from each position in the genome, implemented in ANGSD v0.912 (Korneliussen, Albrechtsen, & Nielsen, 2014). An interspecific genetic distance matrix was constructed using the function “cophenetic.phylo” from the *ape* package (Paradis & Schliep, 2018) with a previously published phylogeny (Kozak et al., 2015). Geographic distance matrices were created by inputting the co-ordinates of collection localities into the function “distm” in the *geosphere* package to calculate the Haversine great-circle-distance between points (Hijmans, 2017).

### Statistical analyses

#### Inter- and intra-specific indicator compounds

We carried out indicator analysis using the *indicspecies* package (Cáceres & Legendre, 2009). Groupings are decided *a priori* (in this case species or sub-species) and compounds are determined which act as indicators of these groups. The best indicators are those which are only found in a single group and all group members possess the compound; such a compound would have an indicator value of 1. We used the function “indicators” to investigate both which single compounds and combinations of compounds best predict group membership. We used the function “pruneindicators” to find the single or combinations of compounds which had the highest indicator values.

#### Variation in chemical profiles

Divergence in chemical profiles across species and populations was estimated with non-metric multidimensional scaling (NMDS) ordination in three dimensions, based on a Bray-Curtis similarity matrix. We used the “metaMDS” function in the *vegan* package version 2.5-1 (Oksanen et al., 2017), and visualised the NMDS using the *ade4* package (Dray & Dufour, 2007).

We assessed the relative importance of relevant factors in driving the variation in chemical profiles with multivariate statistical analyses. These factors included: species identity, geographic region and individual locality. To compare overall variation in chemical composition between groups, we carried out PERMANOVA (permutational multivariate analysis of variance) testing based on a Bray-Curtis distance matrix, using the “adonis2” function in the *vegan* package (Oksanen et al., 2017) with 1000 permutations. We investigated each term in the model sequentially, starting with species identity, the main clustering factor found from visualisation with NMDS, followed by geographic region (Panama *vs* Western Andes *vs* Eastern Andes *vs* Amazon), and finally individual collecting localities. Model goodness of fit was determined was evaluated by Akaike information criterion (AIC). In general, we chose the model with the lowest AIC value, however, if two models were within two AIC of each other we chose the simplest model as the best fit (Table S3). We followed these PERMANOVA tests with *post hoc* pair-wise testing using the function “pairwise.perm.manova” in the *RVAidememoie* package, with Bonferroni correction, to identify which grouping factors were significantly different (Hervé, 2018). We repeated the PERMANOVA analysis within species, in *H. erato* and *H. melpomene*, to investigate fine-scale intraspecific geographic patterns. In the within species analysis we included geographic region (Panama *vs* Western Andes *vs* Eastern Andes *vs* Amazon), and individual collecting localities as the two factors.

One issue with distance-based analyses such as PERMANOVA is that differences in dispersion between groups can be confounded with differences in location (Warton, Wright, & Wang, 2012). To confirm these analyses and account for this issue, we implemented multivariate generalised linear models using the function “ManyGLM” from the *mvabund* package (Wang, Naumann, Wright, & Warton, 2012). We modelled the data using a negative binomial distribution, which we found to be appropriate through examination of residual plots. For interspecific analyses we included species, region, and locality nested within region in the model. For intraspecific analyses we included region and locality nested within region. The “ManyGLM” function fits models to each chemical compound, summing the test statistics to give a multivariate test statistic known as Sum-of-LR. This statistic can be tested for significance using resampling methods. We carried out backwards elimination and compared the fit of models by using the “anova.manyglm” function with a likelihood ratio test (Table S4). We can also determine which compounds are driving between-group differences by looking at the individual contribution of each compound to the Sum-of-LR, with p-values adjusted for multiple testing using the “adjust” option.

### Phylogenetic and geographic distance

Shared ancestry can explain part of the variation in a species’ chemical profile. We tested for a correlation between phylogenetic distance and chemical profile divergence with Mantel tests using the *vegan* package (Oksanen et al., 2017). To investigate the role of geographic distance in chemical profile divergence, we compared geographic and chemical distances matrices with partial Mantel tests. To visualise the species phylogeny (Kozak et al., 2015) we used the “plot.phylo” function from the ape package (Paradis & Schliep, 2018).

### Genomic and chemical distance within species

We calculated intraspecific genetic distances using genome sequences from 11 *H. erato* and 13 *H. melpomene* populations. We visualised genetic distances in two dimensions using MDS with the function “cmdscale”. We tested for a correlation between intraspecific genetic distance and chemical profile divergence with Mantel tests using the *vegan* package (Oksanen et al., 2017). Hybrids between populations of the same species were excluded from this analysis (Table S2). We used partial Mantel tests to investigate the role of geographic distance.

### Co-mimics and similarity of chemical profiles

We used samples of two mimicry rings from two localities, Panama and western Ecuador. *H. melpomene* and *H. erato* form one mimicry ring, whilst *H. cydno* and *H. sapho* form another, with the addition of *H. eleuchia* in western Ecuador (Fig. 1). To test which factors most influenced the variation in chemical profiles we used both PERMANOVA and multivariate GLM. We included species, region, mimicry group, and interactions between these terms in the models.

All statistical analyses were performed with *R* version 3.5.1 (R Core Team, 2018). Figures were made using a palette of colours optimized for colour-blindness (Wong, 2011). Raw sequence data are available through GenBank Short Read Archive (XXX), GCMS data is available through Dryad (XXX), R code and data are available on Open Science Framework (https://osf.io/28yfk/?view_only=c1f7e7a925e74dee84fd2229cbf3f511).

## Results

### Chemical compounds in androconia and genitals

We sampled 252 androconia and 275 genitals across 42 populations of 33 subspecies of seven species, and identified 349 compounds in the genitals and 157 in the androconia. Of the total number of androconial compounds, 38% are fatty acid derivatives, 20% aromatics, 10% terpenoids, 1% macrolides, <1% lactones and 31% unknown or unidentified compounds. Of the genital compounds, 17% are fatty acid derivatives, 7% aromatics, 10% terpenoids, 1% lactones, 12% macrolides and 44% unknown or unidentified compounds. The main difference is that there are more macrolides in the genitals than androconia.

Heliconius species varied considerably in the amount and abundance of compounds (Fig. 2). Between species there was variation in the number of compounds per individual, and the overall amount of compounds detected (Table S5, S6). For the androconia, H. eleuchia had the fewest compounds (13±5), and H. melpomene the most (32±11) (mean ± standard deviation). H. sapho had the lowest total abundance of compounds at 1,300±803 ng, and H. melpomene the most at 7,254±8,242 ng. The species with the fewest genital compounds was H. sapho with 32±7, and the most H. cydno with 102±21. H. sapho also had the lowest total amount of compounds at 6,642±3,975 ng, and H. cydno the highest at 91,167±67,122 ng. These values are within the same order of magnitude as expected from previous work on male sex pheromones in the butterfly Bicyclus anyana (Nieberding et al., 2012; van Bergen, Brakefield, Heuskin, Zwaan, & Nieberding, 2013). Using H. erato as an example, the androconial bouquet is 0.00002%, and genital bouquet 0.0007% of total body weight (Montgomery, Merrill, & Ott, 2016). In general, a higher number of compounds and total abundance of compounds is found in the genitals than in the androconial patches of Heliconius wings.

**Figure 2.**
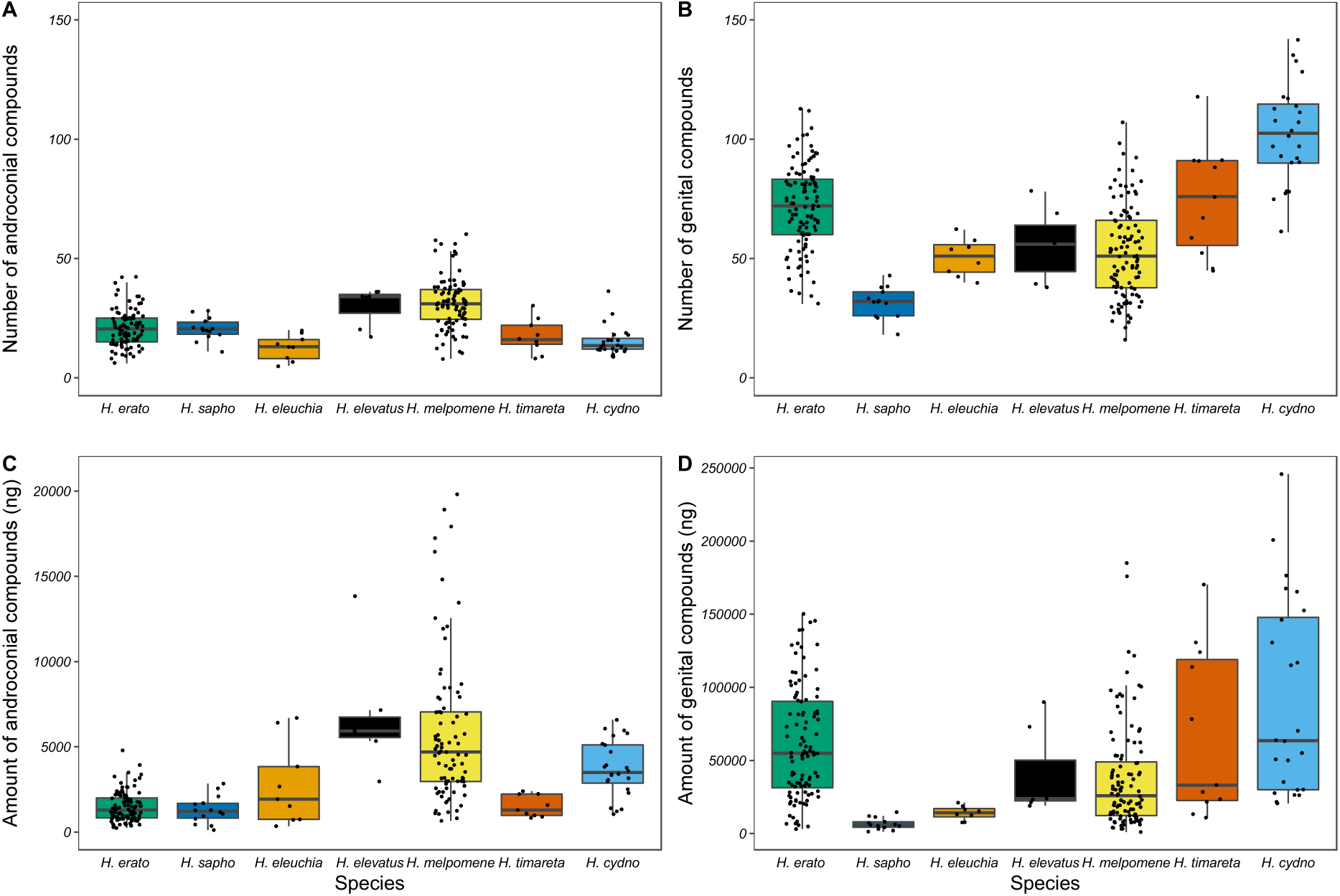
Composition of androconial and genital bouquets across seven *Heliconius* species. Species significantly differ in: (A) number of androconial compounds (ANOVA, F6,245=21.54, p<0.001), (B) number of genital compounds (anova, F6,268=36.15, p<0.001), (C) amount of androconial compounds (ANOVA, F6,245=11.55, p<0.001), (D) amount of genital compounds (anova, F6,268=11.62, p<0.001). Four outlier individuals were removed from C.

### Are there species-specific chemical compounds?

In order to identify candidate species recognition pheromones, we examined our data for species specific compounds using indicator analysis. In most species that we examined there were single androconial compounds that were strong indicators of species identity (Table 1). For example, geranylgeranylacetone was found only in H. erato and was consistently present across samples of this species. Similarily, octadecanal, a compound previously shown to be electrophysiologically active (Byers et al., 2019), was found almost exclusively in H. melpomene (specificity=0.999). H. cydno and H. eleuchia had the weakest indicator scores; in H. cydno because the best indicator compound was not found in all individuals examined (coverage= 0.667), and in H. eleuchia because the best indicator compound was also found in other species (specificity= 0.747). There were similarly species-specific genital compounds in all species except H. sapho and H. timareta, where a combination of two compounds was the best predictor (Table 2). Similar to the androconia, in H. melpomene, the best indicator compound for genitalia has known biological activity, in this case the anti-aphrodisiac, (E)-β-ocimene (Schulz, Estrada, Yildizham, Boppré, & Gilbert, 2008). For H. erato we identified a terpene ester which is only found in H. erato individuals and no other species. Other terpene esters were also almost perfect indicator compounds of H. erato.

**Table 1:**
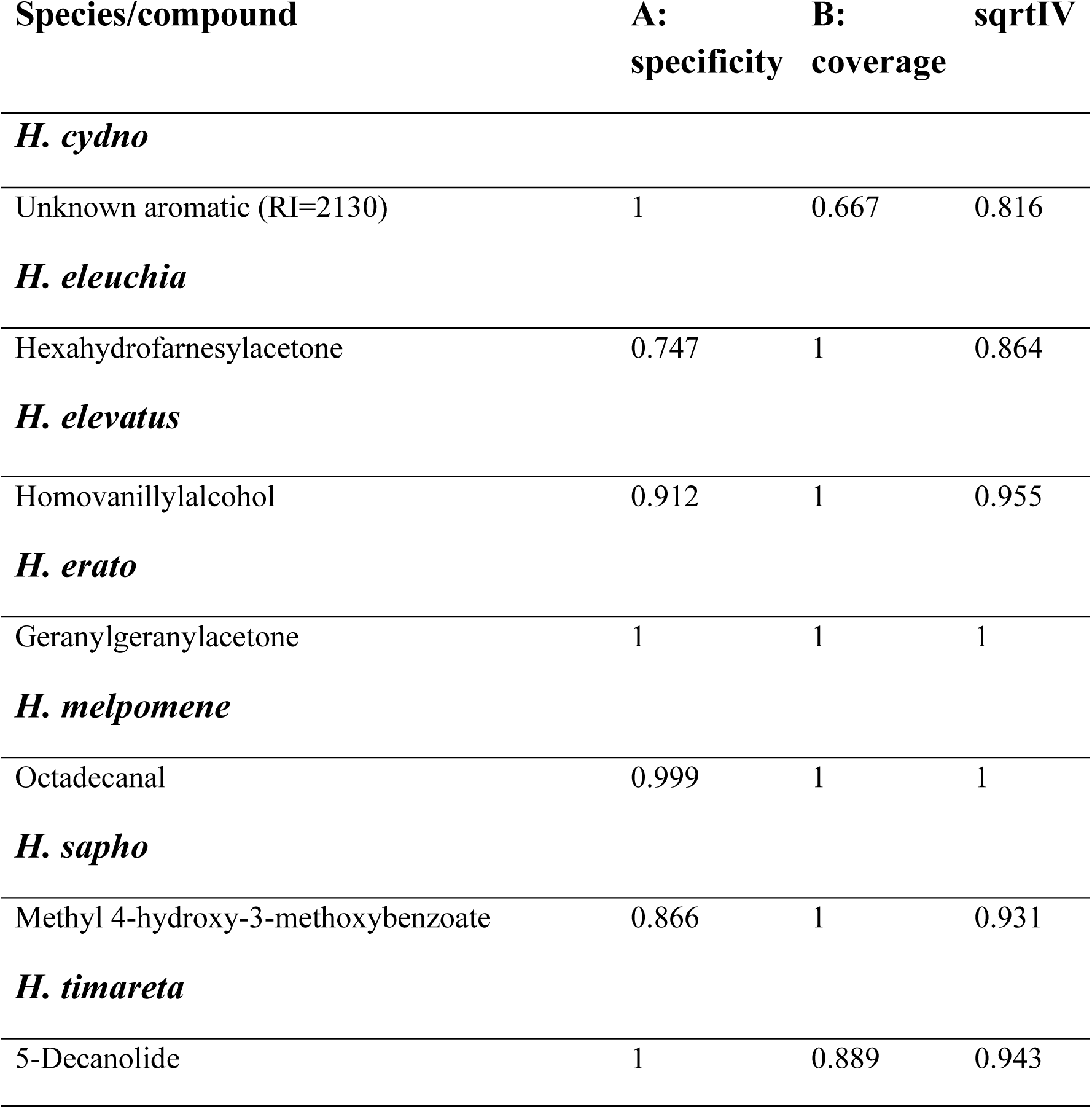
Androconial compounds which are the best indicators of species identity. A is a measure of group specificity of the compounds, B is a measure of group coverage, and sqrtIV is the indicator value which considers both A and B and ranges from 0 (compound not present in any individuals of that species) to 1 (compound only present in that species, and present in all individuals).

**Table 2:** Genital compounds which are the best indicators of species identity. A, B, and sqrtIV as in Table 1.

| **Species/compound** | **A:**<br>**specificity** | **B:**<br>**coverage** | **sqrtIV** |
| --- | --- | --- | --- |
| ***H. cydno*** |  |  |  |
| Unknown ester (RI=1390) | 0.999 | 1 | 0.999 |
| ***H. eleuchia*** |  |  |  |
| Unknown macrolide RI=1878 | 0.969 | 1 | 0.984 |
| ***H. elevatus*** |  |  |  |
| Icosenol | 0.908 | 1 | 0.953 |
| ***H. erato*** |  |  |  |
| Unknown terpene ester (RI=2494) | 1 | 1 | 1 |
| ***H. melpomene*** |  |  |  |
| (*E*)-β-Ocimene | 0.865 | 1 | 0.930 |
| ***H. sapho*** |  |  |  |
| (*Z*)-3-Hexenyl isobutyrate & unknown (RI=1691) | 0.957 | 0.923 | 0.940 |
| ***H. timareta*** |  |  |  |
| Butyl oleate & (*Z*)-9-octadecen-13-olide | 0.915 | 1 | 0.956 |

### What factors affect interspecific variation in chemical profiles?

Our sampling allowed us to investigate how variation in chemical composition is partitioned within and between species, and determine the extent to which chemistry is a species-diagnostic trait. Visualisation of the chemical profiles reveals that individuals mostly group by species for both androconial and genital chemical bouquets (Fig. 3). Species significantly differ in their androconial bouquet, with species identity accounting for 58% of the overall variation in chemical profiles (PERMANOVA, Species, F_6,251_=72.16, p<0.001). All pairwise comparisons of species are significant (Table S7). A further 4% of variation can be explained by region (Amazon/Eastern Andes/Western Andes/Panama), and 3% by locality nested within region (PERMANOVA, Region, F_3,251_=9.96, p<0.001; (Region/Locality), F_8,251_=2.65, p<0.001). Finally, 4% of variation is explained by an interaction between species and region (PERMANOVA, Species*Region F_6,251_=4.82, p<0.001;).

**Figure 3.**
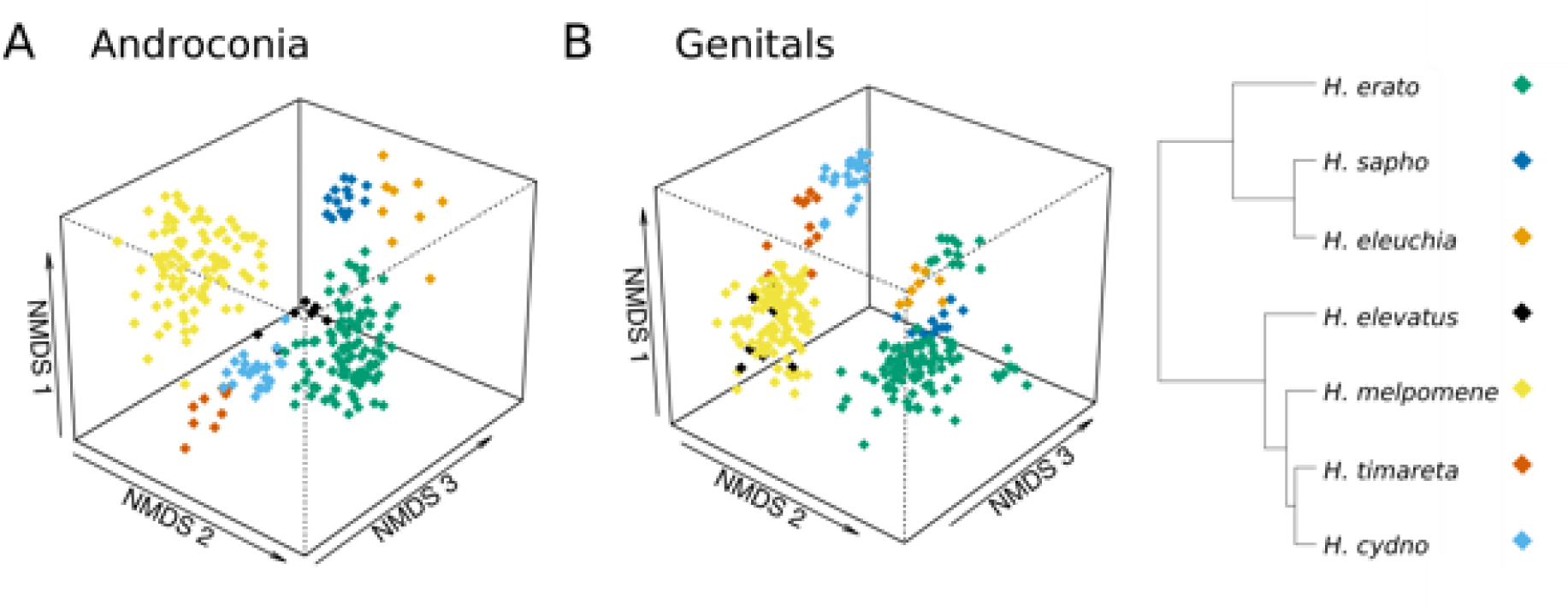
NMDS (non-metric multidimensional scaling) plot illustrating in three dimensions the variation in chemical compounds of male *Heliconius* of different species. A) Androconial compound bouquets differ significantly between species. Stress=0.155. B) Genital bouquets also differ significantly between species. Stress=0.121.

The results were similar for the genital bouquets, with species identity accounting for 51% of the variation in chemical profiles (PERMANOVA, Species, F_6,274_=59.81, p<0.001). All pairwise comparisons are significant apart from H. elevatus and H. melpomene (Table S8). A further 5% of variation can be explained by region (Amazon/Eastern Andes/Western Andes/Panama), and 3% by locality nested within region (PERMANOVA, Region, F_3,274_=12.43, p<0.001; (Region/Locality), F_8,274_=2.92, p<0.001). Finally, 6% of variation is explained by an interaction between species and region (PERMANOVA, Species*Region F_6,274_=6.52, p<0.001). For both androconial and genital chemical profiles, most variation is explained by species identity, rather than geographic location, as confirmed by ManyGLM (Tables S9, S10).

### Does phylogenetic distance explain chemical profile divergence?

Using whole-genome sequence data, we explored the degree to which variation between species can be explained by geographic and genetic distance among the samples. We carried out partial Mantel tests to investigate the correlation between two variables whilst controlling for a third variable. When controlling for geographic distance, genetic divergence is strongly correlated with both androconial and genital chemical divergence (Mantel test, androconia, r=0.7871, p=0.001; genitals, r=0.6936, p=0.001). When controlling for genetic distance, geographic distance is significantly but weakly correlated with androconial and genital chemical divergence (Mantel test, androconia, r=0.072, p=0.001; genitals, r=0.046, p=0.007).

### Do we find subspecies-specific chemical compounds?

We used an indicator analysis to search for compounds unique to specific populations of H. erato and H. melpomene. Most intraspecific differences are due to quantitative rather than qualitative differences between populations, perhaps explaining why many population indicators were weak as they are also found in other regions at different amounts (Table S11, S12). The only exception is H. e. cyrbia (Western Ecuador) that has many genital compounds unique to this region (Table S11).

### What factors affect intraspecific variation in chemical profiles of H. erato and H. melpomene?

We also wanted to determine the sources of variation within species using our broad sampling of populations across the ranges of H. erato and H. melpomene. For H. erato there was a strong grouping of individuals by region (Fig. 4), with 27% of variation in androconial profiles being explained by region and 11% by locality nested within region (PERMANOVA, Region F_3,87_=11.16, p<0.001, Locality F_6,87_=2.35, p<0.001). All four regions are significantly different from each other (Pairwise permutation MANOVAs, p<0.01). For H. erato genital compounds, 37% of variation is explained by region, and 11% by locality nested within region (PERMANOVA, Region F_3,91_=19.01, p<0.001, Locality F_6,91_=2.83, p<0.01). All four regions are significantly different from each other (Pairwise permutation MANOVAs, p<0.05).

**Figure 4.**
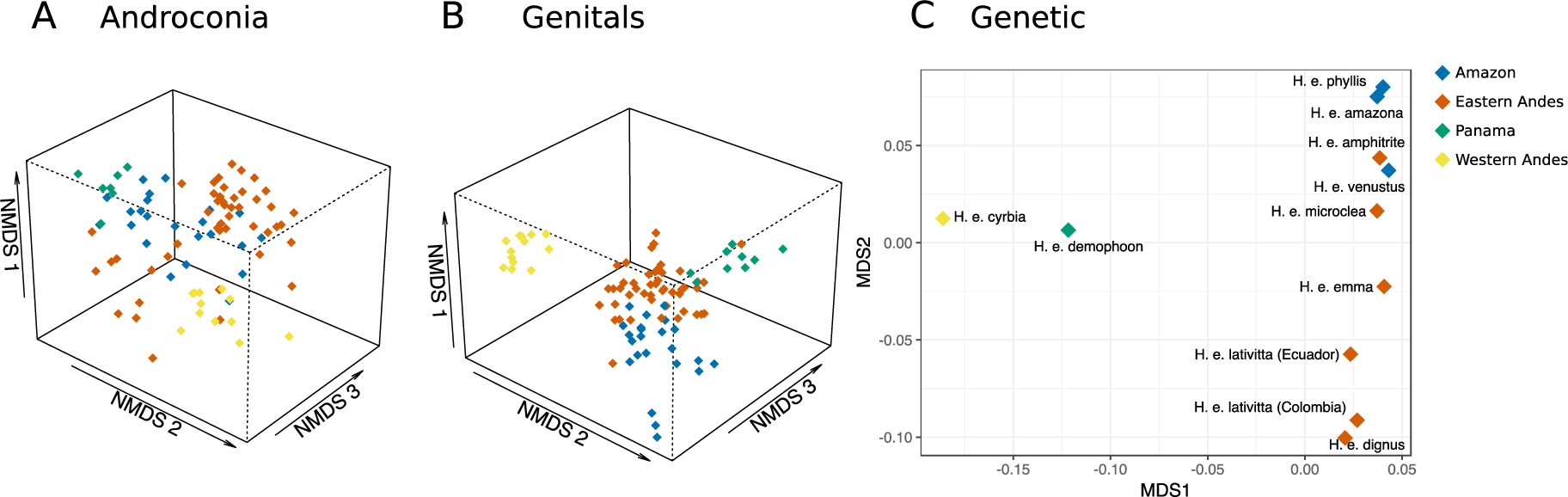
Plots of androconial, genital and genetic distance between *H. erato* populations. A) NMDS (non-metric multidimensional scaling) plot illustrating in three dimensions the variation in androconial chemical compounds. Stress=0.174. B) NMDS plot illustrating in three dimensions the variation in genital chemical compounds. Stress=0.118. C) MDS plot illustrating in two dimensions the genetic distance between populations of *H. erato*.

These geographic differences in chemical profiles are not as strong in H. melpomene (Fig. 5). For H. melpomene androconial compounds, the best model only includes region, not locality, with 18% of variation is explained by region (PERMANOVA, Region F_3,86_=6.73, p<0.01). The West Andes subspecies (H. m. cythera) is not significantly different from either East Andes (multiple populations) or Panama (H. m. rosina) (Pairwise permutation MANOVAs, p=0.072), however, the other comparisons are significantly different (Pairwise permutation MANOVAs, p<0.05). For H. melpomene genital compounds, 20% of variation is explained by region, and 12% by locality nested within region (PERMANOVA, Region F_3,103_=8.91, p<0.001, Locality F_7,103_=2.34, p<0.001). All regions are significantly different from each other (Pairwise permutation MANOVAs, p<0.05), apart from West Andes and Amazon (Pairwise permutation MANOVAs, p=0.120). Both species show variation between geographic locations, with more variance explained by region in H. erato than H. melpomene. These results were confirmed by ManyGLM tests (Tables S13, S14, S15, S16).

**Figure 5.**
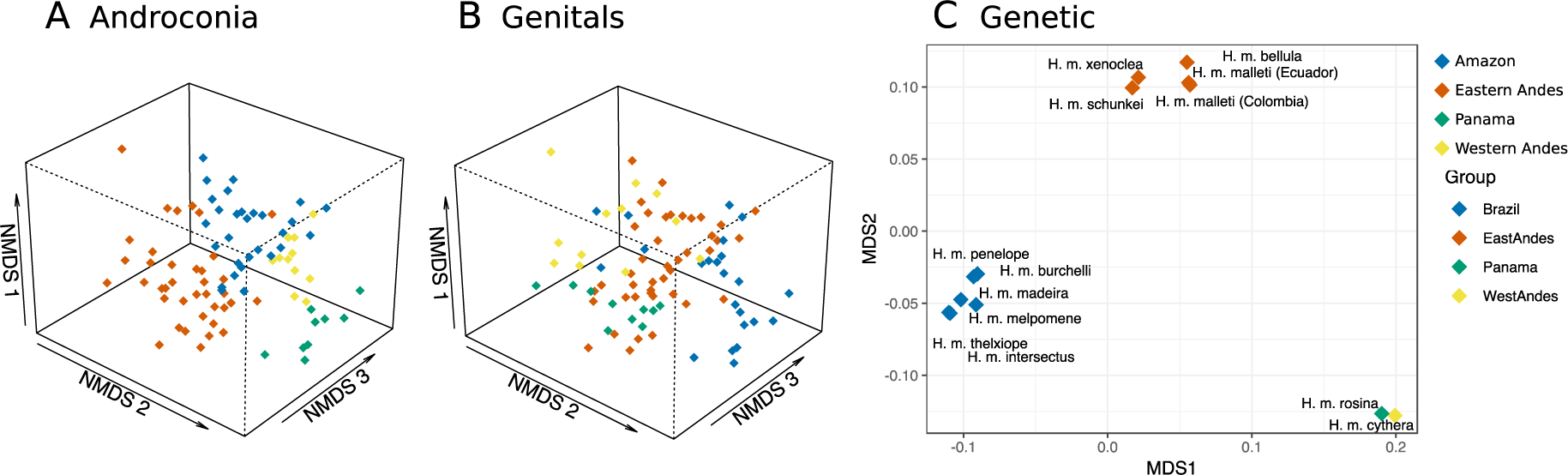
Plots of androconial, genital and genetic distance between *H. melpomene* populations. A) NMDS (non-metric multidimensional scaling) plot illustrating in three dimensions the variation in androconial chemical compounds. Stress=0.151. B) NMDS plot illustrating in three dimensions the variation in genital chemical compounds. Stress=0.161. C) MDS plot illustrating in two dimensions the genetic distance between populations of *H. melpomene*.

### Does genetic distance explain chemical divergence in H. erato and H. melpomene?

In H. erato, chemical distance is positively correlated with genetic distance, when accounting for geographic distance, although this correlation is weak for androconia (partial Mantel test, androconia, R=0.169, p=0.001; genitals, R=0.352, p=0.001). When we account for genetic distance, geographic distance is weakly correlated with androconial chemical distance and not correlated with genital chemical distance (partial Mantel test, androconia, R=0.149, p=0.001; genitals, R=-0.04105, p=0.764). These results were consistent when the analysis was repeated without unknown compounds (supplementary results).

Heliconius melpomene genitals show similar patterns to H. erato, but variation in the androconia is explained by geographic but not genetic distance. When accounting for geography, genetic divergence is not correlated with androconial chemical divergence, and only weakly with genital chemical divergence (partial Mantel test, androconia, R=0.02874, p=0.141, genitals, R=0.1416, p=0.001). When we first consider genetic distance, geographic distance is weakly positively correlated with androconial chemical distance, but not genital chemical distance (partial Mantel test, androconia, R=0.1795, p=0.002; genitals, R=-0.01434, p=0.744).

### Is there evidence for similarity between co-mimics in chemical profile?

We tested for the effect of mimicry ring on chemical profile using individuals collected in Panama and western Ecuador from two mimicry rings (Fig. 6). For androconial chemical profiles, 45% of variation is explained by species (nested within mimicry ring, as each species only belongs to one mimicry ring in our samples), with a further 21% explained by mimicry ring (PERMANOVA, Mimicry ring F_1,85_=54.29, p<0.001; (Mimicry ring/Species) F_3,85_=38.83, p<0.001). A further 2% is explained by region, another 2% by the interaction between region and species nested within mimicry ring, and finally 1% by the interaction between mimicry ring and region (PERMANOVA, Region F_1,85_=4.37, p<0.001; Mimicry ring*Region F_1,85_=2.70, p<0.01; (Mimicry ring/Species)*Region F_2,85_=2.87, p<0.001). We confirmed these results using ManyGLM (Table S17).

**Figure 6.**
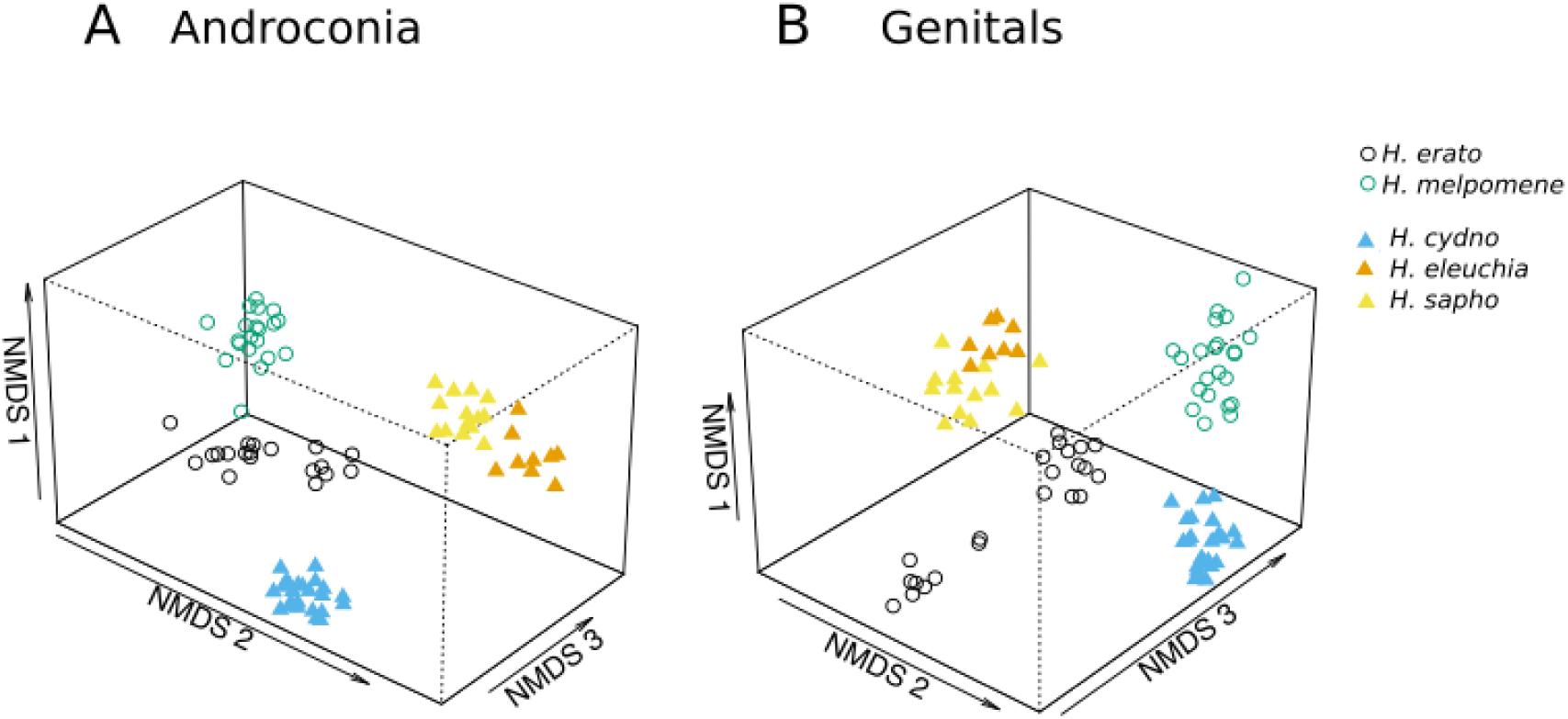
NMDS (non-metric multidimensional scaling) plot illustrating in three dimensions the variation in chemical compounds of male *Heliconius* from Panama and Western Ecuador. *H. erato* and *H. melpomene* are co-mimics (circles), whilst *H. cydno*, *H. eleuchia* and *H. sapho* form a second co-mimicry group (triangles). A) Co-mimicry group contributes 21% of variation in androconial chemical bouquets. Stress=0.098. B) Co-mimicry group contributes 21% of variation in genital chemical bouquets. Stress=0.094.

We found similar trends for genital bouquets (Fig. 6). Again, species (nested within mimicry ring) explained the most variation, 41% of genital bouquet variation, with a further 21% explained by mimicry ring (PERMANOVA, Mimicry ring F_1,88_=56.90, p<0.001; (Mimicry ring/Species) F_3,88_=37.03, p<0.001). A further 3% is explained by region, another 4% by the interaction between region and species nested within mimicry ring, and finally 2% by the interaction between mimicry ring and region (PERMANOVA, Region F_1,88_=9.07, p<0.001; Mimicry ring*Region F_1,88_=4.82, p<0.01; (Mimicry ring/Species)*Region F_2,88_=5.10, p<0.001). We confirmed these results using ManyGLM (Table S18). In both PERMANOVA and in multivariate GLMs, variation in both androconial and genital bouquets is explained mostly by the species of the butterfly, with mimicry ring explaining around half as much variation as species.

All the results described above show a consistent pattern when unidentified compounds were not included in the analysis (supplementary results). Interspecific analyses were also consistent when repeated without populations with a sample of fewer than 5 individuals (this removed 7 populations from androconial analysis and 5 from genital analysis) (supplementary results).

## Discussion

*Heliconius* butterflies represent a continental-scale adaptive radiation (Kozak et al., 2015). Speciation in this group is often associated with divergence in wing colour pattern and pattern variation plays an important role in speciation and mate preference (Jiggins, 2008; Jiggins et al., 2001; Merrill et al., 2015, 2011, 2019; Sánchez et al., 2015). However, one of the surprising findings to emerge from comparative genomic analysis is the wealth of chemosensory genes (Heliconius Genome Consortium, 2012), suggesting that chemical signalling may play an important role in the reproductive biology of the system. To begin to understand the role of chemical signalling in this radiation we have extensively surveyed both inter- and intraspecific variation of *Heliconius*’ androconial and genital chemical profiles across the Neotropics. We find that most of the variation in chemical profile across our samples is explained by species, and we identify key chemicals serving as indicators for each species. Nonetheless, there is also abundant intraspecific variation in chemical profiles. This variation is mainly quantitative in nature, with the exception of *H. erato cyrbia* which has compounds not found in other *H. erato* populations. We also find that co-mimics do exhibit convergence in their chemical profiles, confirming an earlier hypothesis (Mann et al., 2017). Our work sets the stage for further research into the biology and function of chemical profiles, and their role in within and between-species signalling.

When analysing the chemical profiles across a phylogeny it may not be feasible to carry out behavioural tests in every species. We hypothesised that consistent species-specific compounds are likely to be under stabilising selection and therefore to also be biologically important. In *H. melpomene* the compounds identified as indicators for androconia and genitals, octadecanal and (*E*)-β-ocimene respectively, are both known to be biologically active (Byers et al., 2019; Schulz et al., 2008). Combining broad geographic sampling with indicator analysis therefore provides a promising approach to determine potential biologically active compounds in other species, which could then be tested behaviourally. Our analyses have already identified a number of compounds that could now be tested functionally, such as the androconial compound geranylgeranylacetone in *H. erato*.

Chemical profiles are predicted to be highly species-specific if they are involved in species recognition during mating. For instance orchid bee chemical blends, important for mating and species recognition, show high species-specificity, as well as within-species variability, which can be partly explained by geography (Brand et al., 2019; Weber et al., 2016; Zimmermann, Roubik, & Eltz, 2006). We see similar patterns in *Heliconius*, with greater interspecific than intraspecific differences in chemical profiles. Species identity is the best predictor of chemical divergence, with geographic location able to explain some intraspecific differences. One exception to this is *H. elevatus* which does not group separately from its co-mimic *H. melpomene* for genital compounds, despite the fact that these species are not especially closely related in the *Heliconius* phylogeny. Further samples are needed to confirm that this result is not due to the small sample of *H. elevatus* in this study. As in orchid bees, species differences are often consistent across a large geographic range, suggesting that they could be important for reproductive isolation between species (Weber et al., 2016).

We found a correlation between chemical distance and genetic distance. This suggests that neutral evolutionary forces are important in driving chemical bouquet evolution. The correlation between genital chemical distance and genetic distance is a much stronger correlation than previously reported (Catalina Estrada, Schulz, Yildizhan, & Gilbert, 2011), possibly due to the quantitative nature of our data. The strong signal of neutrality suggests that the majority of compounds in the bouquets are neutrally evolving. For example, in the genital bouquet of *H. melpomene*, one compound, (*E*)-β-ocimene, can act by itself as an anti-aphrodisiac, with other components of the bouquet thought to moderate its evaporation rate (Schulz et al., 2008). In the future, focusing on the evolutionary patterns of only biologically active compounds, rather than the entire bouquet, may disentangle the processes involved in the evolution of these profiles.

*Heliconius erato* and *H. melpomene* both exhibit extensive colour pattern variation across their geographic range (Sheppard, Turner, Brown K. S., Benson W. W., & Singer M. C., 1985) and these populations also differ in their androconial and genital bouquets. Whilst traditionally predicted to be under stabilising selection, intraspecific variation between populations in chemical profiles has been documented in other Lepidoptera (Bacquet et al., 2016; Groot et al., 2009; Huang et al., 1998; McElfresh & Millar, 2001; Takanashi et al., 2005). Chemical divergence in putative male sex pheromones between populations of *Bicyclus anynana* is reported to be as large as differences between *Bicyclus* species, and is greater than predicted by genetic divergence (Bacquet et al., 2016). This is in contrast to what we find here, where interspecific differences are much greater than intraspecific ones.

Interestingly, *Heliconius erato cyrbia* produces many unique genital compounds and is also the most genetically divergent *H. erato* population in our study, suggesting that genetic drift is important for the evolution of chemical profiles in *Heliconius*. Across all *H. erato* populations we find a correlation between chemical distance and genetic distance, which is weaker for androconial bouquets. In *H. melpomene*, genetic distance is also weakly correlated with genital chemical divergence. These correlations suggest that some of the geographic variation between populations could be neutral, with stochastic processes important for bouquet evolution in *Heliconius*. In contrast, androconial chemical variation in *H. melpomene* is better explained by geographic distance. This does not match our expectations of neutrality and might imply that other evolutionary forces are important for chemical profile evolution in *H. melpomene*. One such factor might be larval host plant use, which affects compound production (Darragh et al., 2019) and varies across the geographic range of *H. melpomene* (Benson, 1978; Benson, Brown, & Gilbert, 1975).

*Heliconius* butterflies are an excellent example of visual mimicry, with different species converging on the same warning colour patterns (Merrill et al., 2015; Sheppard et al., 1985; Sherratt, 2008). It has been suggested that chemical compounds could also contribute to mimicry between species (Dettner & Liepert, 1994; Mann et al., 2017). In this study, we find evidence that individuals within co-mimicry groups have more similar chemical profiles than expected. Most known examples of chemical mimicry come from systems of deception, for example, mimicry of ant alarm pheromones by rove beetles to avoid predation, rather than mimicry of aposematic warning signals (Dettner & Liepert, 1994; Stoeffler, Maier, Tolasch, & Steidle, 2007; Vereecken & McNeil, 2010). We suggest that in *Heliconius* different components of the bouquet could be important for chemical mimicry and species recognition, reducing conflict between these selection pressures.

Convergence of genital bouquets between co-mimics could be due to the anti-aphrodisiac function of these compounds (Gilbert, 1976; Schulz et al., 2008). Anti-aphrodisiac compounds are transferred from males to females during mating to deter future matings from other males. Convergence in wing pattern between co-mimics could result in harassment not only by conspecific but also heterospecific males (C. Estrada & Jiggins, 2008). The use of the same anti-aphrodisiac by co-mimics could combat interspecific attraction by deterring males of both species, as highlighted by the production of (*E*)-β-ocimene by *H. erato* and *H. melpomene*

Even compounds which are not involved in chemical mimicry could play a role in predator deterrence. Genital compounds were originally suggested to form part of the anti-predation signal (Eltringham, 1925). We detected 2-sec-butyl-3-methoxypyrazine in the genitals of *H. melpomene*, *H. cydno* and *H. timareta*, and 2-isobutyl-3-methoxypyrazine in the genitals of *H. melpomene* and *H. cydno*, both compounds known to deter predators in the wood tiger moth (Burdfield-Steel, Pakkanen, Rojas, Galarza, & Mappes, 2018; Rojas et al., 2018, 2017; Rojas, Mappes, & Burdfield-Steel, 2019). More generally, methoxypyrazines act as warning odours in other insects (e.g. Lepidoptera, Rothschild, Moore & Brown, 1984; fireflies, Vencl et al., 2016), effective against avian predators (Guilford, Nicol, Rothschild, & Moore, 1987). Further investigation will be required to determine if odours of *Heliconius* butterflies act as anti-predation signals.

Overall, our study reveals strong species differences in bouquets and the presence of species-specific compounds, as well as intraspecific variation. A pattern of species-specificity alongside intraspecific variation geographically could be the result of a balance between stabilising selection towards a species stereotype, sexual selection promoting diversity, and geographic segregation alongside selection and drift. A challenge for the field is the feasibility of testing for the biological relevance of hundreds of compounds in many species, but we hope that our innovative analysis will stimulate not only further targeted functional studies of putatively important compounds, but also large chemical profile surveys in other study systems of evolutionary interest.

## Acknowledgements

KD and GMK were supported by Natural Research Council Doctoral Training Partnership (NE/L002507/1) and KD was additionally supported by a Smithsonian Tropical Research Institute Short Term Fellowship KMK was funded by a Smithsonian Institution Fellowship. CRM was supported by a Smithsonian Tropical Research Institute Short Term Fellowship and Predoctoral Fellowship. This study was financed in part by the Coordenação de Aperfeiçoamento de Pessoal de Nível Superior – Brasil (CAPES) – Finance Code 001 (MSc Scholarship for CMEF). CMEF was also funded by a Smithonsian Tropical Research institute Short Term Fellowship. CS was funded by COLCIENCIAS (FP44842-5-2017). ML was supported by research funds from Universidad del Rosario. KJRPB and CDJ were supported by the European Research Council (339873 SpeciationGenetics). RMM was supported by an DFG Emmy Noether fellowship. WOM was supported by the Smithsonian Tropical Research Institute and NSF (DEB 1257689). SS thanks the Deutsche Forschungsgemeinschaft (DFG) for support through grant Schu984/12-1. We thank Michel Cast for photos of *H. melpomene madeira* and *H. m. intersectus* (https://cliniquevetodax.com/Heliconius/). The authors have no conflicts of interest to declare.

## Author contributions

Kathy Darragh, Gabriela Montejo-Kovacevich, Krzysztof Kozak, Colin Morrison, Owen McMillan, Clarisse Figueiredo, Jonathan Ready, Camilo Salazar, Mauricio Linares and Chris Jiggins collected samples. Kathy Darragh analysed the data. Stefan Schulz assisted with the chemical analysis. Gabriela Montejo-Kovacevich created the intraspecific genetic distance matrices. Richard Merrill, Owen McMillan, Kelsey Byers and Chris Jiggins contributed to project design. All authors contributed to manuscript revisions.

## Data availability

Raw sequence data are available through GenBank Short Read Archive (XXX), GCMS data is available through Dryad (XXX), R code and data are available on Open Science Framework (https://osf.io/28yfk/?view_only=c1f7e7a925e74dee84fd2229cbf3f511).

