## Supplementary File for "Species specificity and intraspecific variation in the chemical profiles of *Heliconius* butterflies across a large geographic range"

### Supplementary results

*Does removing unidentified compounds or poorly sampled populations affect models of interspecific variation in chemical profiles?*

We repeated the interspecific analysis without unidentified compounds. When repeated without unidentified compounds, species still significantly differ in their androconial bouquet, with species identity accounting for 60% of the overall variation in chemical profiles (PERMANOVA, Species, $F_{6,251}$=77.33, p<0.001). A further 4% of variation can be explained by region (Amazon/Eastern Andes/Western Andes/Panama), and 3% by locality nested within region (PERMANOVA, Region, $F_{3,251}$=9.96, p<0.001; (Region/Locality), $F_{8,251}$=2.49, p<0.001). Finally, 4% of variation is explained by an interaction between species and region (PERMANOVA, Species*Region $F_{6,251}$=4.77, p<0.001;). Results were also similar for genital bouquets, with species identity still explaining the highest amount ofvariation, accounting for 44% of the variation in chemical profiles (PERMANOVA, Species, $F_{6,274}$=45.44, p<0.001). A further 6% of variation can be explained by region (Amazon/Eastern Andes/Western Andes/Panama), and 4% by locality nested within region (PERMANOVA, Region, $F_{3,274}$=12.74, p<0.001; (Region/Locality), $F_{8,274}$=2.85, p<0.001). Finally, 6% of variation is explained by an interaction between species and region (PERMANOVA, Species*Region $F_{6,274}$=6.32, p<0.001).

We also repeated the interspecific analyses removing populations with fewer than 5 individuals. Again, similar to removing unidentified compounds, and the full dataset, species identity accounts for 58% of overall variation in chemical profiles, with a further 5% explained by region, 3% by locality nested within region, and 4% by an interaction between species and region ( PERMANOVA, Species, $F_{5,227}$=78.26, p<0.001; Region, $F_{3,227}$=10.19, p<0.001; (Region/Locality), $F_{7,227}$=2.98, p<0.001; Species*Region $F_{4,227}$=6.92, p<0.001;). Results were also consistent for genital bouquets, with species identity still explaining the highest amount ofvariation, accounting for 51% of the variation in chemical profiles, with a further 5% explained by region, 4% by locality nested within region, and 5% by an interaction between species and region (PERMANOVA, Species, $F_{5,255}=69.49$, $p<0.001$; Region, $F_{3,255}=12.37$, $p<0.001$; (Region/Locality), $F_{8,255}=3.05$, $p<0.001$; Species*Region $F_{4,255}=9.13$, $p<0.001$).

*Does removing unidentified compounds or poorly sampled populations affect correlations between divergence in chemical profile with genetic and geographic distance?*

Correlations with genetic and geographic distances were also consistent with results including all compounds. When controlling for geographic distance, genetic divergence is strongly correlated with both androconial and genital chemical divergence (Mantel test, androconia, r=0.7897, p=0.001; genitals, r=0.5203, p=0.001). When controlling for genetic distance, geographic distance is significantly but weakly correlated with chemical divergence (Mantel test, androconia, r=0.06739, p=0.002; genitals, r=0.059, p=0.003).

Removing populations with fewer than 5 individuals also gave consistent results. When controlling for geographic distance, genetic and chemical divergence remain strongly correlated (Mantel test, androconia, r=0.7978, p=0.001; genitals, r=0.71, p=0.001). Again, when controlling for genetic distance, geographic distance is significantly but weakly correlated with androconial chemical divergence (Mantel test: r=0.082, p=0.001), however, in this case the weak correlation with genital chemical divergence is not significant (Mantel test: r=-0.0416, p=0.996).

*Does removing unidentified compounds affect models of intraspecific variation in chemical profiles?*

Individuals of *H. erato* still strongly group by region when unidentified compounds are removed from the analysis, with 27% of variation in androconial profiles being explained by region and 11% by locality nested within region (PERMANOVA, Region $F_{3,87}=11.49$, p<0.001, Locality $F_{6,87}$=2.30, p<0.001). Again, this is similar for *H. erato* genital compounds. Region explains 35% of variation, and 8% is explained by locality nested within region (PERMANOVA, Region $F_{3,91}$=16.76, p<0.001, Locality $F_{6,91}$=1.98, p<0.01).

Again, we found consistent results without unidentified compounds for *H. melpomene.* The same amount of variation was explained in models with and without unidentified compounds included, with region explaining 18% of variation in androconial compounds (PERMANOVA, Region $F_{3,86}$=6.05, p<0.01). For *H. melpomene* genital compounds, 20% of variation is explained by region, 12% by locality nested within region, as in models with all compounds included (PERMANOVA, Region $F_{3,103}$=9.05, p<0.001, Locality $F_{7,103}$=2.34, p<0.001).

*Does removing unidentified compounds affect correlations between intraspecific chemical divergence with genetic and geographic distance?*

Results were again very similar without unidentified compounds included in the analysis. In *H. erato* both androconial and genital chemical distance are positively correlated with genetic distance, even when accounting for geographic distance (partial Mantel test, androconia, R=0.154, p=0.001; genitals, R=0.3116, p=0.001). When unidentified compounds are removed, we still find a weak positive correlation between geographic distance and androconial, but not genital distance, accounting for genetic distance (partial Mantel test, androconia, R=0.1527, p=0.001; genitals, R=-0.0773, p=0.949).

For *H. melpomene*, genital bouquet divergence, but not androconial bouquet divergence, is correlated with genetic distance when accounting for geography (partial Mantel test, androconia, R=0.02602, p=0.187, genitals, R=0.1362, p=0.001). When we first consider genetic distance, geographic distance is not positively correlated with genital chemical distance, however, it is positively correlated with androconial chemical distance (partial Mantel test, androconia, R=0.1729, p=0.002; genitals, R=-0.01255, p=0.732). These results are consistent with tests including all compounds.

*Does removing unidentified compounds change models of similarity between co-mimics?*

The effect of mimicry ring on chemical profile is consistent when unidentified compounds are removed. For androconial chemical profiles, 45% of variation is explained by species (nested within mimicry ring, as each species only belongs to one mimicry ring in our samples), with a further 21% explained by mimicry ring (PERMANOVA, Mimicry ring $F_{1,85}=55.24$, $p<0.001$; (Mimicry ring/Species) $F_{3,85}=39.48$, $p<0.001$). A further 2% is explained by region, another 2% by the interaction between region and species nested within mimicry ring, and finally 1% by the interaction between mimicry ring and region (PERMANOVA, Region $F_{1,85}=3.99$, $p<0.001$; Mimicry ring*Region $F_{1,85}=2.42$, $p<0.01$; (Mimicry ring/Species)*Region $F_{2,85}=2.59$, $p<0.001$).

We found similar trends for genital bouquets when unidentified compounds were removed. Again, species (nested within mimicry ring) explained the most variation, 36% of genital bouquet variation, with a further 22% explained by mimicry ring (PERMANOVA, Mimicry ring $F_{1,88}=56.28$, $p<0.001$; (Mimicry ring/Species) $F_{3,88}=30.51$, $p<0.001$). A further 5% is explained by region, another 4% by the interaction between region and species nested within mimicry ring, and finally 2% by the interaction between mimicry ring and region (PERMANOVA, Region $F_{1,88}=11.67$, $p<0.001$; Mimicry ring*Region $F_{1,88}=5.36$, $p<0.01$; (Mimicry ring/Species)*Region $F_{2,88}=5.25$, $p<0.001$).

**Table S1: Number of androconial and genital samples collected for each locality**

| Locality | Species | No. androconial samples | No. genital samples |
|---|---|---|---|
| Panama | *H. cydno* | 12 | 12 |
| Panama | *H. erato* | 7 | 9 |
| Panama | *H. melpomene* | 9 | 10 |
| Panama | *H. sapho* | 12 | 11 |
| Colombia (highlands) | *H. erato* | 10 | 9 |
| Colombia (highlands) | *H. melpomene* | 13 | 14 |
| Colombia (highlands) | *H. timareta* | 7 | 8 |
| Colombia (lowlands) | *H. erato* | 12 | 13 |
| Colombia (lowlands) | *H. melpomene* | 7 | 8 |
| Ecuador (east) | *H. elevatus* | 2 | 2 |
| Ecuador (east) | *H. erato* | 10 | 11 |
| Ecuador (east) | *H. melpomene* | 11 | 13 |
| Ecuador (east) | *H. timareta* | 2 | 3 |
| Ecuador (west) | *H. cydno* | 12 | 12 |
| Ecuador (west) | *H. eleuchia* | 9 | 8 |
| Ecuador (west) | *H. erato* | 12 | 13 |
| Ecuador (west) | *H. melpomene* | 11 | 12 |
| Ecuador (west) | *H. sapho* | 2 | 2 |
| Peru (central) | *H. erato* | 7 | 6 |
| Peru (central) | *H. melpomene* | 3 | 4 |
| Peru (south) | *H. erato* | 9 | 9 |
| Peru (south) | *H. melpomene* | 3 | 3 |
| Brazil (Coastal Para) | *H. melpomene* | 8 | 8 |
| Brazil (Mato Grosso) | *H. erato* | 12 | 12 |
| Brazil (Mato Grosso) | *H. melpomene* | 10 | 15 |
| Brazil (North Para) | *H. erato* | 5 | 6 |
| Brazil (North Para) | *H. melpomene* | 13 | 15 |
| Brazil (Rondonia) | *H. elevatus* | 3 | 3 |
| Brazil (Rondonia) | *H. erato* | 10 | 11 |
| Brazil (South Para) | *H. elevatus* | 2 | 2 |
| Brazil (South Para) | *H. erato* | 4 | 5 |
| Brazil (South Para) | *H. melpomene* | 3 | 6 |

**Table S2: Genome samples of *H. erato* and *H. melpomene* races including number of androconial and genital samples included in analysis for each race**

| Taxon name | No. androconial samples | No. genital samples | SampleID | Sex | Longitude | Latitude | Accession |
|---|---|---|---|---|---|---|---|
| *H. melpomene bellula* | 13 | 14 | CAM040049 | M | -76.683 | 1.217 | |
| *H. melpomene malleti* | 7 | 8 | ERR1307111 | M | -75.669 | 1.814 | SAMEA3723397 |
| *H. melpomene malleti* | 11 | 13 | 0ERR350930 | M | -77.668 | -1.061 | SAMEA2240083 |
| *H. melpomene cythera* | 11 | 12 | 14N015 | M | -78.853 | 0.185 | |
| *H. melpomene burchelli* | 3 | 5 | burchelliSR281 | M | -56.404 | -13.814 | |
| *H. melpomene intersectus* | 1 | 1 | intersectusKK291 | | | | |
| *H. melpomene madeira* | 2 | 5 | madeiraSR391 | M | -54.847 | -4.066 | |
| *H. melpomene melpomene* | 13 | 15 | melpomeneSR178 | M | -54.626 | -1.937 | |
| *H. melpomene penelope* | 4 | 7 | penelopeSR358 | M | -57.706 | -13.691 | |
| *H. melpomene thelxiope* | 7 | 7 | thelxiopeKK288 | | | | |
| *H. melpomene schunkei* | 3 | 3 | schunkeiKK544 | M | -70.768 | -13.204 | |
| *H. melpomene xenoclea* | 3 | 4 | xenocleaKK309 | M | -75.407 | -11.0354 | |
| *H. melpomene rosina* | 9 | 10 | ERR2298205 | M | -79.659 | 9.076 | SAMEA104585083 |
| *H. erato dignus* | 10 | 9 | CAM040113 | M | -76.690 | 1.214 | |
| *H. erato lativitta* | 12 | 13 | CAM040160 | M | -76.409 | 0.956 | |
| *H. erato cyrbia* | 12 | 13 | CAM040545 | M | -78.770 | 0.151 | |
| *H. erato lativitta* | 10 | 11 | CAM041030 | M | -77.702 | -1.059 | |
| *H. erato amazona* | 4 | 5 | amazonaSR122 | | | | |
| *H. erato phyllis* | 10 | 8 | phyllisSR230 | M | -55.440 | -10.891 | |
| *H. erato venustus* | 9 | 10 | venustusSR314 | M | -60.297 | -12.806 | |
| *H. erato amphitrite* | 9 | 9 | amphitriteKK464 | M | -72.656 | -12.955 | |
| *H. erato emma* | 1 | 1 | emmaKK402 | M | -74.935 | -10.298 | |
| *H. erato microclea* | 4 | 4 | microcleaKK338 | M | -75.419 | -11.055 | |
| *H. erato demophoon* | 7 | 9 | SRR4032093 | M | 79.715 | -9.129 | SAMN05224182 |

**Table S3. Model selection table for PERMANOVA models based on AIC scores. If two models were within two AIC points of each other, we chose the simpler model as the most parsimonious. Best fit models are highlighted in bold.**

| Model | Residual Sum of Squares | DF | AIC |
|---|---|---|---|
| Interspecific androconia | | | |
| Chemical profile ~ Species + Region + (Region/Locality) + Species*Region + Species*(Region/Locality) | 26.436 | 31 | 889.2356 |
| **Chemical profile ~ Species + Region + (Region/Locality) + Species*Region** | 28.185 | 23 | 889.3774 |
| Chemical profile ~ Species + Region + (Region/Locality) | 31.760 | 17 | 907.4649 |
| Chemical profile ~ Species + Region | 34.377 | 9 | 911.4244 |
| Chemical profile ~ Species | 38.07 | 6 | 931.1351 |
| Interspecific genitals | | | |
| Chemical profile ~ Species + Region + (Region/Locality) + Species*Region + Species*(Region/Locality) | 31.887 | 31 | 1016.104 |
| **Chemical profile ~ Species + Region + (Region/Locality) + Species*Region** | **33.748** | **23** | **1015.703** |
| Chemical profile ~ Species + Region + (Region/Locality) | 39.008 | 17 | 1043.535 |
| Chemical profile ~ Species + Region | 42.151 | 9 | 1048.847 |
| Chemical profile ~ Species | 47.165 | 6 | 1073.751 |
| *H. erato* androconia | | | |
| **Chemical profile ~ Region + (Region/Locality)** | 8.79 | 9 | 211.2867 |
| Chemical profile ~ Region | 10.3793 | 3 | 213.9033 |
| *H. erato* genitals | | | |
| **Chemical profile ~ Region + (Region/Locality)** | 9.1105 | 9 | 223.2669 |
| Chemical profile ~ Region | 10.9982 | 3 | 228.5917 |
| *H. melpomene* androconia | | | |

| | | | |
|---|---|---|---|
| Chemical profile ~ Region + (Region/Locality) | 9.8523 | 11 | 223.0307 |
| **Chemical profile ~ Region** | 11.7698 | 3 | 222.502 |
| *H. melpomene* genitals | | | |
| **Chemical profile ~ Region + (Region/Locality)** | 12.3471 | 10 | 283.3958 |
| Chemical profile ~ Region | 14.5178 | 3 | 286.2391 |
| Mimicry ring androconia | | | |
| **Chemical profile ~ Mimicry ring + (Mimicry ring/Species) + Region + Mimicry*Region + (Mimicry/Species)*Region** | 9.660 | 8 | 213.0481 |
| Chemical profile ~ Mimicry ring + (Mimicry ring/Species) + Region + Mimicry*Region | 10.379 | 6 | 215.2235 |
| Chemical profile ~ Mimicry ring + (Mimicry ring/Species) + Region | 10.718 | 5 | 215.9874 |
| Chemical profile ~ Mimicry ring + (Mimicry ring/Species) | 11.144 | 4 | 217.3353 |
| Chemical profile ~ Mimicry ring | 25.882 | 1 | 283.8048 |
| Mimicry ring genitals | | | |
| **Chemical profile ~ Mimicry ring + (Mimicry ring/Species) + Region + Mimicry*Region + (Mimicry/Species)*Region** | 9.699 | 8 | 220.2101 |
| Chemical profile ~ Mimicry ring + (Mimicry ring/Species) + Region + Mimicry*Region | 10.935 | 6 | 226.8874 |
| Chemical profile ~ Mimicry ring + (Mimicry ring/Species) + Region | 11.520 | 5 | 229.5224 |
| Chemical profile ~ Mimicry ring + (Mimicry ring/Species) | 12.525 | 4 | 234.9698 |
| Chemical profile ~ Mimicry ring | 26.088 | 1 | 294.2722 |

**Table S4. Analysis of deviance model selection table for multivariate generalised linear models based on likelihood ratio tests. Each model is compared to the model above it. Best fit models are highlighted in bold.**

| Model | ΔDeviance | Residual DF | p-value |
|---|---|---|---|
| Interspecific androconia | | | |
| **Chemical profile ~ Species + Region + (Region/Locality) + Species*Region + Species*(Region/Locality)** | | 220 | |
| Chemical profile ~ Species + Region + (Region/Locality) + Species*Region | -740.7 | 228 | 0.001 |
| Interspecific genitals | | | |
| Chemical profile ~ Species + Region + (Region/Locality) + Species*Region + Species*(Region/Locality) | | 243 | |
| Chemical profile ~ Species + Region + (Region/Locality) + Species*Region | -1854 | 251 | 0.063 |
| **Chemical profile ~ Species + Region + (Region/Locality)** | -2787 | 257 | 0.134 |
| Chemical profile ~ Species + Region | -6481 | 265 | 0.026 |
| *H. erato* androconia | | | |
| **Chemical profile ~ Region + (Region/Locality)** | | 78 | |
| Chemical profile ~ Region | -1091 | 84 | 0.001 |
| *H. erato* genitals | | | |
| **Chemical profile ~ Region + (Region/Locality)** | | 82 | |
| Chemical profile ~ Region | -2721 | 88 | 0.001 |
| *H. melpomene* androconia | | | |
| **Chemical profile ~ Region + (Region/Locality)** | | 75 | |
| Chemical profile ~ Region | -1582 | 83 | 0.001 |
| *H. melpomene* genitals | | | |
| **Chemical profile ~ Region + (Region/Locality)** | | 93 | |

| | | | |
|---|---|---|---|
| Chemical profile ~ Region | -1777 | 100 | 0.001 |
| Mimicry ring androconia | | | |
| **Chemical profile ~ Mimicry ring + (Mimicry ring/Species) + Region + Mimicry*Region + (Mimicry/Species)*Region** | | 77 | |
| Chemical profile ~ Mimicry ring + (Mimicry ring/Species) + Region + Mimicry*Region | -156.5 | 79 | 0.001 |
| Mimicry ring genitals | | | |
| **Chemical profile ~ Mimicry ring + (Mimicry ring/Species) + Region + Mimicry*Region + (Mimicry/Species)*Region** | | 80 | |
| Chemical profile ~ Mimicry ring + (Mimicry ring/Species) + Region + Mimicry*Region | -567.4 | 82 | 0.001 |

**Table S5: Androconial compounds identified in at least half of all individuals from any population. The gas chromatographic retention index is reported for each compound. Mean amounts (ng) in each species are reported. (Attached excel file)**

**Table S6 – Genital compounds identified in at least half of all individuals from any population. The gas chromatographic retention index is reported for each compound. Mean amounts (ng) in each species are reported. (Attached excel file)**

**Table S7: Pairwise PERMANOVA p-values of between species androconial comparisons using Bonferroni correction. All values were significant at 0.05 significance level.**

| | *H. cydno* | *H. eleuchia* | *H. elevatus* | *H. erato* | *H. melpomene* | *H. sapho* |
|---|---|---|---|---|---|---|
| *H. eleuchia* | 0.021 | | | | | |
| *H. elevatus* | 0.042 | 0.021 | | | | |
| *H. erato* | 0.021 | 0.021 | 0.021 | | | |
| *H. melpomene* | 0.021 | 0.021 | 0.021 | 0.021 | | |
| *H. sapho* | 0.021 | 0.021 | 0.021 | 0.021 | 0.021 | |
| *H. timareta* | 0.021 | 0.021 | 0.021 | 0.021 | 0.021 | 0.021 |

**Table S8: Pairwise PERMANOVA p-values of between species genital comparisons using Bonferroni correction. Significant results are highlighted in bold.**

| | *H. cydno* | *H. eleuchia* | *H. elevatus* | *H. erato* | *H. melpomene* | *H. sapho* |
|---|---|---|---|---|---|---|
| *H. eleuchia* | **0.021** | | | | | |
| *H. elevatus* | **0.021** | **0.021** | | | | |
| *H. erato* | **0.021** | **0.021** | **0.021** | | | |
| *H. melpomene* | **0.021** | **0.021** | 0.273 | **0.021** | | |
| *H. sapho* | **0.021** | **0.042** | **0.021** | **0.021** | **0.021** | |
| *H. timareta* | **0.021** | **0.021** | **0.021** | **0.021** | **0.021** | **0.021** |

**Table S9: Summary of androconial chemical bouquet analysis of all species using the ManyGLM approach including all significant explanatory variables. The ten compounds which contribute the most to the deviance explained by a variable are listed for each variable in descending order of contribution. Compounds highlighted with * were also identified by an indicator analysis.**

| Parameter | Residual DF | DF | Deviance | p-val | Compounds |
|---|---|---|---|---|---|
| Species | 245 | 6 | 10944 | 0.001 | Geranylgeranylacetone*, syringaldehyde, methyloctadecanal (RI=2076), icosanal, octadecanal*, (*Z*)-11-icosenal, henicosane, methyloctadecanal (RI=2064), unknown RI=1396, 1-hexadecanol |
| Region | 242 | 3 | 3717 | 0.001 | Henicosane, tricosane, unknown ester RI=1188, unknown RI=2133, tetracosane, napthalene[1], unknown RI=1366, unknown RI=2277, pentacosane |
| (Region/Locality) | 231 | 11 | 2826 | 0.001 | Hexadecadien-15-olide, unknown RI=1915, unknown hydrocarbon RI=1962, 1-hexadecanol, napthalene[1], nonanal, (*Z*)-13-docosenal, methyl 4-hydroxy-3-methoxybenzoate, (*Z*)-β-ocimene, unknown RI=1184 |
| Species*Region | 222 | 9 | 896 | 0.005 | Napthalene, methyl salicylate, henicosane, 1-octadecanol, mellein, dihydroactinidiolide, 1-hexadecanol, octadecanal, (*Z*)-13-docosen-1-ol, tricosane |
| Species*(Region/Locality) | 220 | 21 | 741 | 0.001 | 1-Hexadecanol, pentacosane, 1-octadecanol, methyl salicylate, henicosane |

[1]Naphthalene is a known flower volatile, but can also be introduced by contamination. Our blank samples never contained naphthalene, indicating the butterfly origin in our study.

**Table S10: Summary of genital chemical bouquet analysis of all species using the ManyGLM approach including all significant explanatory variables. The ten compounds which contribute the most to the deviance explained by a variable are listed for each variable. Compounds highlighted with * were also identified by an indicator analysis.**

| Parameter | Residual DF | DF | Deviance | p-val | Compounds |
|---|---|---|---|---|---|
| Species | 268 | 6 | 27587 | 0.001 | Unknown terpene ester RI=2494*, unknown terpene ester RI=2139, henicosane, unknown pentyl ester RI=2033, unknown terpene ester 2435, unknown aromatic RI=1299, unknown terpene RI=2755, benzyl cyanide, unknown sesterterpene hydrocarbon RI=2370, (*E*)-β-ocimene* |
| Region | 265 | 3 | 8965 | 0.001 | Unknown RI=2840, 7,8-dihydro-β-ionone, benzyl cyanide, unknown aromatic ester RI=2511, 2-phenylethyl dodecanoate, unknown triterpene RI=2891, unknown aromatic ester RI=2718, hexadecane, unknown RI=1076, 3-undecanone |
| (Region/Locality) | 257 | 11 | 6431 | 0.016 | (*Z*)-β-Ocimene, unknown RI=1915, unknown hydrocarbon RI=1750, 18-octadecanolide, henicosene (2068), 19-methylicosyl acetate, napthalene, hexadecanoic acid, 3-undecanone, unknown terpene ester RI=2310 |

**Table S11: Androconial and genital compounds which are the best indicators of different geographic groups of *H. erato*. A is a measure of group specificity of the compounds, B is a measure of group coverage, and sqrtIV is the indicator value which considers both A and B and ranges from 0 (compound not present in any individuals of that species) to 1 (compound only present in that species, and present in all individuals).**

| ***Wings*** <br> ***Amazon*** | **A: specificity** | **B: coverage** | **sqrtIV** |
|---|---|---|---|
| Napthalene | 0.787 | 0.870 | 0.827 |
| ***East Andes*** | | | |
| 1-Hexadecanol & mellein | 0.802 | 0.957 | 0.876 |
| ***West Andes*** | | | |
| Unknown RI=1704 | 0.950 | 0.833 | 0.890 |
| ***Panama*** | | | |
| Benzylacetate | 1 | 1 | 1 |
| Unknown ester RI=1188 | 1 | 1 | 1 |
| ***Genitals*** | | | |
| ***Amazon*** | | | |
| Napthalene | 0.851 | 0.957 | 0.902 |
| ***East Andes*** | | | |
| Unknown triterpene RI=2891 | 0.846 | 0.978 | 0.910 |
| ***West*** | | | |
| Unknown RI=1833 | 1 | 1 | 1 |
| Unknown RI=1970 | 1 | 1 | 1 |
| 2-Phenylethyl decanoate | 1 | 1 | 1 |
| Unknown terpene ester RI=2120 | 1 | 1 | 1 |
| 2-Phenylethyl dodecanoate | 1 | 1 | 1 |
| Unknown RI=2258 | 1 | 1 | 1 |
| 2-Phenylethyl tetradecenoate | 1 | 1 | 1 |
| Unknown aromatic ester RI=2511 | 1 | 1 | 1 |
| Unknown aromatic ester RI=2718 | 1 | 1 | 1 |
| Unknown RI=2734 | 1 | 1 | 1 |
| ***Panama*** | | | |
| Benzyl acetate | 1 | 1 | 1 |
| Unknown ester RI=1188 | 1 | 1 | 1 |
| Pentyl/isopentyl 3-methylbutyrate | 1 | 1 | 1 |

**Table S12: Androconial and genital compounds which are the best indicators of different geographic groups of *H. melpomene*. A is a measure of group specificity of the compounds, B is a measure of group coverage, and sqrtIV is the indicator value which considers both A and B and ranges from 0 (compound not present in any individuals of that species) to 1 (compound only present in that species, and present in all individuals).**

| ***Wings*** <br> ***Amazon*** | **A: specificity** | **B: coverage** | **sqrtIV** |
|---|---|---|---|
| Alkene or alcohol (RI=2127) & henicosane | 0.871 | 0.966 | 0.917 |
| ***East Andes*** | | | |
| (*Z*)-13-Docosenal & henicosane | 0.962 | 0.895 | 0.928 |
| ***West Andes*** | | | |
| Unknown RI=1766 | 1 | 0.727 | 0.852 |
| ***Panama*** | | | |
| Nonanoic acid | 0.816 | 1 | 0.903 |
| ***Genitals*** <br> ***Amazon*** | | | |
| 14-Tetradecanolide | 0.915 | 0.975 | 0.945 |
| ***East Andes*** | | | |
| 7,8-Dihydro-β-ionone | 1 | 0.881 | 0.939 |
| ***West*** | | | |
| Hexyl octadecenoate (RI=2621) | 0.826 | 1 | 0.909 |
| ***Panama*** | | | |
| 2-*sec*-Butyl-3-methoxypyrazine | 0.495 | 1 | 0.704 |

**Table S13: Summary of androconial chemical bouquet analysis of *H. erato* using the ManyGLM approach including all significant explanatory variables. The ten compounds which contribute the most to the deviance explained by a variable are listed for each variable. Compounds highlighted with * were also identified by an indicator analysis.**

| Parameter | Residual DF | DF | Deviance | p-val | Compounds |
|---|---|---|---|---|---|
| Region | 84 | 3 | 1275 | 0.001 | Pentacosane, unknown ester RI=1188*, benzyl acetate*, unknown macrolide RI=1714, tetracosane, unknown RI=1366, pentyl/isopentyl 3-ethylbutyrate RI=1145, mellein*, hexadecadien-15-olide, heptadecene, |
| (Region/Locality) | 78 | 9 | 1091 | 0.001 | Hexadecadien-15-olide, napthalene*, unknown RI=1915, unknown hydrocarbon RI=1962, tetracosane, unknown RI=1184, unknown macrolide RI=1714, pentacosane, unknown RI=1444, unknown RI=1424 |

**Table S14: Summary of genital chemical bouquet analysis of *H. erato* using the ManyGLM approach including all significant explanatory variables. The ten compounds which contribute the most to the deviance explained by a variable are listed for each variable. Compounds highlighted with * were also identified by an indicator analysis.**

| Parameter | Residual DF | DF | Deviance | p-val | Compounds |
|---|---|---|---|---|---|
| Region | 88 | 3 | 6142 | 0.001 | 3-Undecanone, unknown sesterterpene RI=2636*, unknown RI=2840, unknown tripterpene RI=2891*, unknown aromatic ester RI=2511*, 2-phenylethyl dodecanoate*, unknown RI=2451, unknown terpene ester RI=2435, unknown aromatic ester RI=2718*, benzyl cyanide, |
| (Region/Locality) | 82 | 9 | 2721 | 0.001 | Napthalene*, hexadecanoic acid, unknown diterpene RI=2205, unknown ester hexanoate RI=1565, 18-octadecanolide, unknown RI=2279, unknown macrolide RI=1714, unknown RI=1424, unknown amide RI=2157, icosanal |

**Table S15: Summary of androconial chemical bouquet analysis of *H. melpomene* using the ManyGLM approach including all significant explanatory variables. The ten compounds which contribute the most to the deviance explained by a variable are listed for each variable.**

| Parameter | Residual DF | DF | Deviance | p-val | Compounds |
|---|---|---|---|---|---|
| Region | 83 | 3 | 1848 | 0.001 | Henicosane*, tricosane, methyl 3,4-dimethoxybenzoate, homovanillyl alcohol, (*Z*)-13-docosen-1-ol*, 11-icosenol, icosenol, napthalene, unknown aromatic RI=1738, unknown alkene or alcohol RI=2127* |
| (Region/Locality) | 75 | 11 | 1582 | 0.001 | Unknown RI=2133, nonanal, 1-octadecanol, (*Z*)-13-docosenal, (*Z*)-9-octadecenal, Unknown RI=1915, (*Z*)-16-methyl-9-octadecenol, unknown RI=2112, unknown RI=1638, tricosene RI=2072 |

**Table S16: Summary of genital chemical bouquet analysis of *H. melpomene* using the ManyGLM approach including all significant explanatory variables. The ten compounds which contribute the most to the deviance explained by a variable are listed for each variable. Compounds highlighted with * were also identified by an indicator analysis.**

| Parameter | Residual DF | DF | Deviance | p-val | Compounds |
|---|---|---|---|---|---|
| Region | 100 | 3 | 2281 | 0.001 | 7,8-Dihydro-β-ionone*, 12-dodecanolide, 2-*sec*-butyl-3-methoxypyrazine, napthalene, unknown RI=1704, 2-methoxy-3-isobutylpyrazine*, (*Z*)-β-ocimene, unknown RI=1607, 11-dodecanolide, (*E*)-β-ocimene |
| (Region/Locality) | 93 | 10 | 1777 | 0.001 | 12-Dodecanolide, unknown hydrocarbon RI=1750, unknown RI=1915, benzyl salicylate, hexenyl octadecatrienoate & (Z)-3-hexenyl octadecenoate, 11-methylpentacosane, unknown RI=2891, 14-tetradecanolide*, unknown sesquiterpene RI=1902, 11-dodecanolide |

**Table S17: Summary of androconial chemical bouquet analysis of *H. melpomene, H. erato, H. sapho, H. cydno* and *H. eleuchia* from western Ecuador and Panama using the ManyGLM approach including all significant explanatory variables. The ten compounds which contribute the most to the deviance explained by a variable are listed for each variable (less than ten are listed for the interaction terms as these are the only significant compounds).**

| Parameter | Residual DF | DF | Deviance | p-val | Compounds |
|---|---|---|---|---|---|
| Mimicry ring | 84 | 1 | 1527 | 0.001 | 1-Octadecanol, 1-hexadecanol, methyloctadecanal RI=2076, gernaylgeranylacetone, hexadecadien-15-olide, methyloctadecanal RI=2064, icosane, (*E*)-phytol, hexahydrofarnesyl acetone, henicosane |
| Region | 83 | 1 | 463 | 0.001 | Unknown RI=1188, unknown RI=1366, benzyl acetate, unknown RI=1915, 11-icosenol, unknown aromatic RI=1757, decanal, tricosene, unknown hydrocarbon RI=1962, napthalene |
| (Mimicry ring/Species) | 78 | 5 | 3001 | 0.001 | Syringaldehyde, henicosane, octadecanal, icosanal, geranylgeranylacetone, hexahydrofarnesyl acetone, (*E*)-phytol, tricosane, methyl 4-hydroxy-3-methoxybenzoate, icosane |
| Mimicry ring*Region | 77 | 1 | 202 | 0.001 | Methyl salicylate, (*Z*)-13-docosen-1ol, octadecanal, 19-methylicosyl acetate, 1-octadecanol, icosenol |
| (Mimicry ring/Species) *Region | 77 | 3 | 157 | 0.001 | Napthalene, hexadecanal, 1-hexadecanol, methyl salicylate, nonanal, mellein |

**Table S18: Summary of genital chemical bouquet analysis of *H. melpomene*, *H. erato*, *H. sapho*, *H. cydno* and *H. eleuchia* from western Ecuador and Panama using the ManyGLM approach including all significant explanatory variables. The ten compounds which contribute the most to the deviance explained by a variable are listed for each variable.**

| Parameter | Residual DF | DF | Deviance | p-val | Compounds |
|---|---|---|---|---|---|
| Mimicry ring | 87 | 1 | 4577 | 0.001 | (*Z*)-3-Hexyl hexanoate (branched?), 16-hexadecanolide, (Z)-3-hexenyl hexanoate, (*E*)-β-ocimene, unknown ester RI=1392, (*Z*9,*E*11)- 9,11-octadecadien -13-olide, butyl oleate, butyl hexadecanoate, (*Z*)-9-octadecen-11-olide, unknown hexenyl or cyclo pentyl ester RI=1775 |
| Region | 86 | 1 | 1581 | 0.001 | Unknown ester RI=1188, unknown RI=2840, unknown RI=2670, unknown RI=1970, benzyl acetate, hexadecanoic acid, phenylethanol, unknown compound RI=2451, 2-phenylethyl dodecanoate |
| (Mimicry ring/Species) | 81 | 5 | 10034 | 0.001 | Benzyl cyanide, henicosane, unknown terpene ester RI=2310, unknown RI=1396, hexyl (*E*)-2-hexenoate, (*Z*)-9-octadecen-11-olide, (*Z*9,*E*11)- 9,11-octadecadien -13-olide, unknown terpene ester RI=2494, isopropyl octadecadienolate |
| Mimicry ring*Region | 80 | 1 | 897 | 0.001 | Unknown RI=2657, unknown aromatic RI=1299, 2-phenylethyl decanoate, benzyl hexadecanoate, unknown sesterterpene RI=2636, unknown RI=2166, hexyl-3-methyl butyrate, (*Z*)-3-hexenyl isobutyrate, unknown RI=2864, heptacosene |
| (Mimicry ring/Species) *Region | 80 | 3 | 567 | 0.011 | Unknown sesterterpene RI=2636, hexyl-3-methyl butyrate, (*Z*)-3-hexenyl isobutyrate, henicosene RI=2072, (Z)-3-hexenyl hexadecanoate, (*E*)-β-ocimene, dihydroedulan II, hexyl octadecenoate & (Z)-3-hexenyl octadecenoate, hexenyl octadecatrienoate & (Z)-3-hexenyl octadecatrienoate, napthalene |
